# Insulin-Regulated Actin Dynamics is Disrupted in a Human Keratinocyte Model of Hailey Hailey Disease

**DOI:** 10.64898/2025.12.15.694174

**Authors:** Ruby Gupta, Akash Chinchole, Ngozi P. Paul, Mrinal K. Sarkar, Johann E. Gudjonsson, Ajay Verma, Rajini Rao

**Affiliations:** Department of Physiology, Pharmacology & Therapeutics, Johns Hopkins University School of Medicine, 725 N. Wolfe St, Baltimore MD 21205; Department of Dermatology, Taubman Medical Research Institute, University of Michigan, Ann Arbor, MI 48109; Formation Venture Engineering Foundry, 47 Fox Run Road, Topsfield MA

**Keywords:** ATP2C1, secretory pathway calcium ATPase, wound healing, lamellipodia, insulin, Rac1

## Abstract

Hailey Hailey Disease (HHD) is an autosomal dominant cutaneous disorder caused by mutations in *ATP2C1*, the gene encoding the Golgi/secretory pathway Ca^2+^-ATPase SPCA1. Characterized by suprabasal acantholysis and intertriginous blistering of the skin, HHD treatment focuses on managing symptoms as there is no cure. Challenges to targeted therapy are due to the lack of facile and reliable models, both human and rodent, for mechanistic studies. Here we validate and characterize CRISPR/Cas9 mediated single and bi-allelic *ATP2C1* knockouts in immortalized human hTERT keratinocytes. Whereas SPCA1 expression, Golgi morphology and Golgi Ca^2+^ accumulation were proportionately affected in heterozygous and homozygous *ATP2C1* null mutants as expected, both single and double allelic mutants showed near complete loss of cadherins associated with desmosomal and adherens junctions. HHD is characterized by poor wound healing and impaired keratinocyte migration. We show that SPCA1 is required for dynamic reorganization of actin cytoskeleton in keratinocyte spreading. We identified an insulin activated PI3K-AKT-Rac1 signaling pathway required for lamellipodia formation and keratinocyte spreading, defective in SPCA1 mutants. Transgenic expression of hSPCA1 or treatment with CDN1163, a small molecule Ca^2+^-ATPase agonist, restored defective phenotypes in the HHD model, paving the way for future therapeutic approaches to treat this disorder.

## Introduction

Hailey-Hailey disease (HHD) is an autosomal dominant genodermatosis caused by mutations in *ATP2C1* affecting approximately 1 in 50,000 people [1, 2]. The gene encodes the <u>S</u>ecretory <u>P</u>athway <u>C</u>a^2+^-<u>A</u>TPase isoform 1 protein, SPCA1, which localizes to the Golgi apparatus of the keratinocyte. Patients develop painful, blistering skin lesions that effect the neck, armpit, chest, skin folds and groin followed by secondary fungal infections of the affected areas [3]. In keratinocytes, defective biogenesis and trafficking of desmosomal components causes acantholysis in suprabasal layers of skin resulting in a characteristic “dilapidated brick wall” appearance [4]. Despite the known molecular etiology, no FDA-approved treatment exists and patients are symptomatically treated with topical or oral antibiotics, corticosteroids, cyclosporine or methotrexate [5]. There are emerging case studies showing relief with the opioid antagonist naltrexone, although results are variable [6]. Laser ablation and surgery can provide a measure of relief to some [7]. Unfortunately, these treatments eventually fail for most patients, and the disease is essentially incurable and debilitating.

The lack of targeted therapy is partly due to the absence of animal models: murine *Atp2c1* homozygous knockout is embryonic lethal, and the heterozygous model fails to exhibit skin acantholysis typical of patients [8, 9]. Species-specific differences in epidermal calcium gradients and keratinization make human-based in vitro systems essential for disease modeling and therapeutic testing [10]. Cell culture models for in vitro characterization of HHD mutations typically involve overexpression [11, 12] or have non-mammalian origins [13, 14] that do not mimic keratinocyte disease etiology. A commonly used keratinocyte model, the HaCaT cell line is aneuploid and has altered transcriptional profiles of genes crucial for the skin’s barrier function [15]. Although biopsy-derived HHD patient samples are invaluable [16, 17], they are difficult to obtain, short lived and show inter-patient variability. Therefore, a key technical requirement is the engineering of human keratinocyte cell lines that will accurately and reliably model HHD.

While immunohistochemical and electron microscopic examination of HHD biopsies have revealed internalization and loss of junctional desmosomal-keratin filament complexes, alterations in the adherens junction and actin cytoskeleton have also been observed in the form of prominent microvilli-like protrusions that appear to compensate for the loss of desmosomes [18]. Actin fibers were observed to be preferentially arranged in parallel bundles beneath the cell membrane in lesional areas from patient skin [18, 19]. Consistent with the ATP-dependent function of SPCA1 to regulate cytoplasmic and Golgi Ca^2+^ levels, Ca^2+^ and ATP-dependent actin organization was found to be defective in HHD keratinocytes [20].

Given these observations, we were interested in understanding the role of SPCA1 in actin cytoskeleton mediated cell spreading and migration. The ability of keratinocytes to spread and change shape dynamically is vital for maintaining skin integrity and facilitating the rapid repair of the essential skin barrier after injury. Wound healing or re-epithelialization requires keratinocytes at the wound edge to change shape, flatten and extend lamellipodia and filopodia which allows them to move laterally across the wound bed [21]. Critical to this process are signaling pathways involving growth factors and small GTPases that regulate the protrusion and cell-matrix attachment of the leading edge, with concomitant retraction and detachment of the lagging edge of the cell resulting in coordinated and directional movement that ensures efficient wound closure. HHD is characterized by poor wound healing [3] and downregulation of *ATP2C1* impairs keratinocyte migration in a scratch wound assay [22]. Here, we use CRISPR/Cas9 mediated single and double allele knockouts of *ATP2C1* in human hTERT/KerCT keratinocytes to show a role for Ca^2+^ homeostasis in cell spreading. Further we uncover an insulin regulated signaling pathway leading to Rac1 GTPase activation required for lamellipodia formation and keratinocyte spreading that is defective in the HHD model. These defects could be reversed by modest transgenic expression of *ATP2C1* or activation of the related ER-localized SERCA pump, pointing to therapeutic opportunities in gene therapy and treatment with small molecule agonists of Ca^2+^-ATPases.

## Results

### Construction and validation of human keratinocytes with single or bi-allelic *ATP2C1* gene deletions

We used hTERT/Ker-CT keratinocytes that were generated by transducing human foreskin keratinocytes with human telomerase reverse transcriptase and mouse Cdk4 [23, 24]. These cells have been karyotyped and shown to differentiate normally in both monolayer and 3D organotypic cultures [25]. They have been successfully used to model psoriasis and atopic dermatitis and validated for drug screening purposes [25, 26]. Single and double allele knockouts of the *ATP2C1* gene were generated using CRISPR/Cas9 and validated as described in Methods. **Fig. 1** compares SPCA1 expression, Golgi localization and Golgi Ca^2+^ accumulation in keratinocytes. hTERT keratinocytes express SPCA1 that co-localizes in the perinuclear region with the Golgi marker GM130 as seen by immunofluorescence microscopy (**Fig. 1a**) and by Western blotting (**Fig. 1b-c**). Whereas the bi-allelic *ATP2C1* knockout had no discernable SPCA1 expression, the mono-allelic knockout showed partial expression that co-localizes weakly with GM130. In control keratinocytes, the Golgi appeared as contiguous ribbons, seen by 3D reconstruction of GM130 immunofluorescence (**Fig. S1a**). In contrast, the Golgi ribbons appeared fragmented in both *ATP2C1* mutant lines (**Fig. S1b-c**), as previously reported in SPCA1 knockdown or knockout models [8, 27, 28].

**Figure 1.**
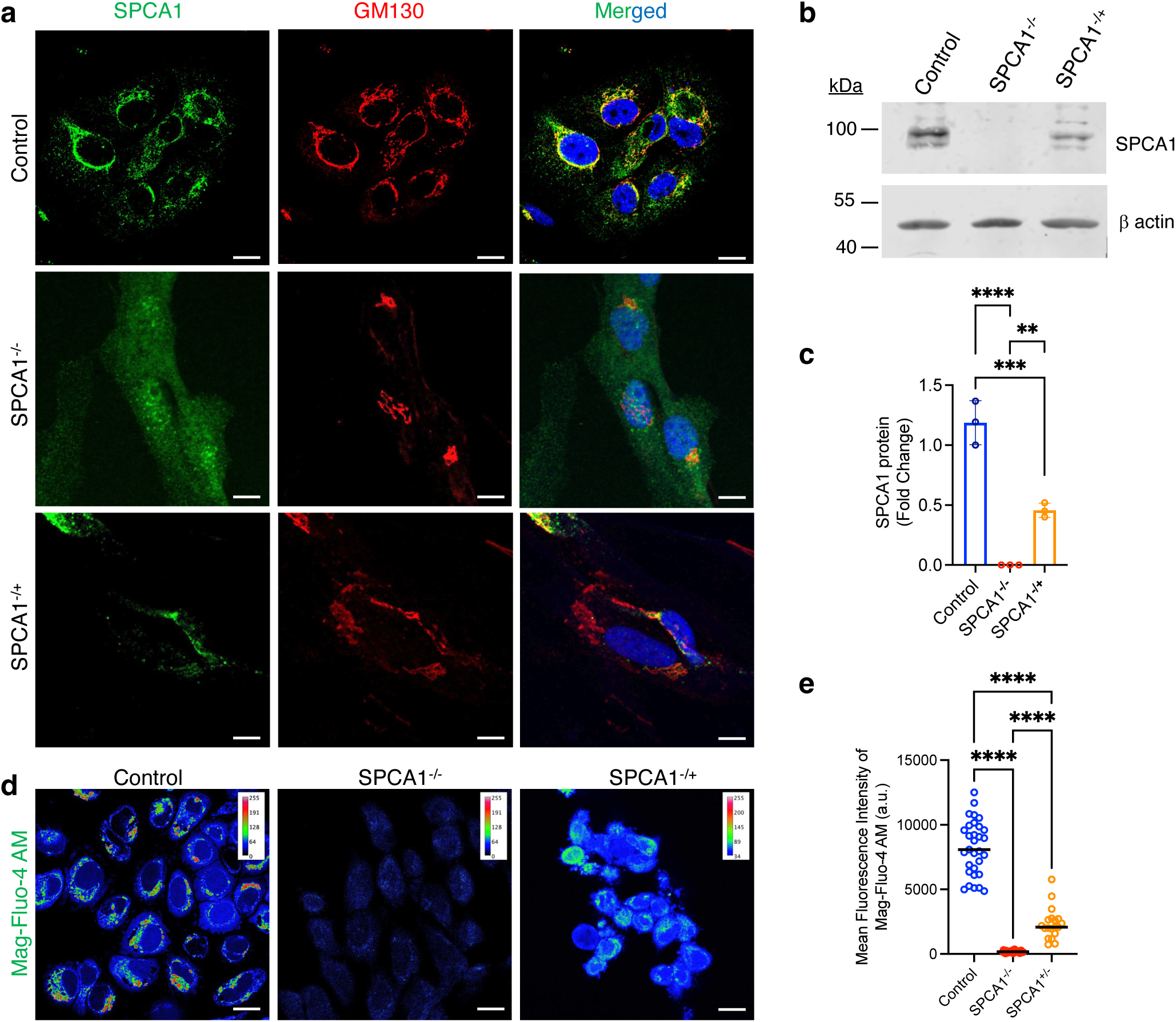
Validation of hTERT keratinocytes with single or bi-allelic *ATP2C1* gene deletions. **(a)** Representative immunofluorescence images of Golgi (GM130) and SPCA1 in three keratinocyte lines: hTERT (Control), single knockout (SPCA1^-/+^) and biallelic knockout (SPCA2^-/-^). Magnification, 63x. Scale bar = 10 μm. **(b)** Representative Western blot and **(c)** densitometric quantification of total levels of SPCA1 relative to β-actin loading control. SPCA1 was reduced on average by 55% in single allelic knockout whereas the bi-allelic knockout showed no detectable expression. **(d)** Representative fluorescence images with **(e)** quantification of Mag-Fluo-4 AM in keratinocytes representing Golgi calcium stores. Magnification, 40x. Scale bar = 20 μm. All graphs represent mean ± SD for n=3 replicates per group. Data were evaluated using Ordinary One-way ANOVA analysis. *\*\*P<0.01; ***P<0.001; ****P<0.0001*.

The accumulation of Golgi Ca^2+^ by the secretory pathway Ca^2+^-ATPase is critical for protein sorting, processing and quality control [9, 29, 30]. Recently, the Mg^2+^/Ca^2+^ selective fluorescent probe MagFluo-4 AM has been used to report on Golgi Ca^2+^ [31, 32]. We observed strong perinuclear fluorescence in hTERT keratinocytes loaded with MagFluo-4 which was absent or sharply reduced in the SPCA1^-/-^ and SPCA1^-/+^ keratinocytes, respectively (**Fig. 1d-e**) consistent with a loss of Golgi Ca^2+^ stores in the SPCA1 mutants. Taken together, these data confirm the partial and complete knockout of *ATP2C1* in the keratinocyte cell lines as expected.

### SPCA1 mutant keratinocytes recapitulate HHD junctional phenotypes

The diagnostic characteristic of HHD is acantholytic dyskeratosis caused by disruption of keratinocyte junctions chiefly due to abnormalities in the desmosome-keratin filament structure [18]. Whereas biopsies from healthy skin showed strong peripheral staining for desmosomal proteins (e.g., Dsg1), lesional areas from HHD patients showed complete dissolution of the desmosomal complex and diffuse staining of Dsg1 in the cytoplasm [33, 34]. We observed near complete absence of Dsg1, Dsg2 and Dsg3 in SPCA1^-/-^ keratinocytes and weak peripheral staining in SPCA1^-/+^ keratinocytes relative to hTERT control (**Fig. 2a; Fig. S2a-b**). To confirm these observations, we treated hTERT keratinocytes with the SPCA1-selective inhibitor bisphenol [35]. **Fig. S2c-d** shows dose-dependent loss of desmoglein proteins by immunofluorescence. Abnormalities in the adherens junctions in lesional skin have also been reported. We observed near complete loss of E-cadherin in both single and bi-allelic knockout keratinocyte lines, as seen by immunofluorescence and Western blotting (**Fig. 2b; Fig. S2e-f**).

**Figure 2.**
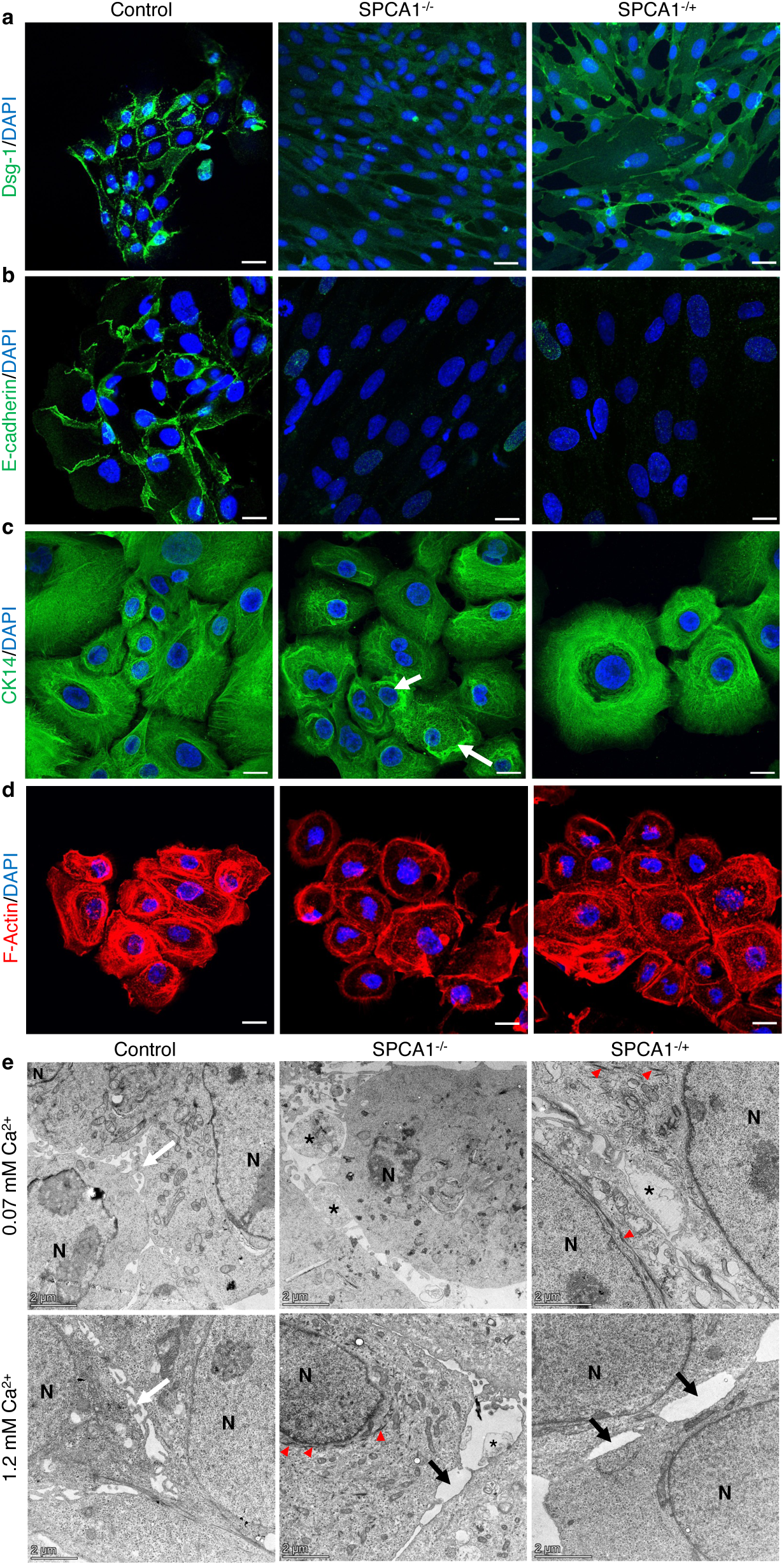
Loss of SPCA1 impairs cell junctions and cytoskeletal integrity. Representative immunofluorescence images of **(a)** Dsg-1 showing strong peripheral localization in control keratinocytes, but markedly reduced or absent staining in SPCA1^-/-^and SPCA1^+/−^ cells. Magnification, 20x. Scale bar = 20 μm; **(b)** E-cadherin immunofluorescence showing robust expression at cell junctions in control keratinocytes but near absence in SPCA1 mutants. Magnification, 20x. Scale bar = 20 μm; **(c)** CK-14 immunofluorescence revealing prominent perinuclear keratin bundles in SPCA1^−/−^ keratinocytes, with fewer aggregates in SPCA1^+/-^ cells. Magnification, 40x. Scale bar = 10 μm; and **(d)** F-actin labeled with fluorescent phalloidin demonstrates abnormal peripheral actin accumulation and cytoplasmic actin-rich structures in SPCA1 mutant cells. Magnification, 40x. Scale bar = 10 μm. **(e)** Transmission electron micrographs showing intercellular gaps (black arrows), membrane blebs (asterisks) and keratin bundles (red arrowheads) in SPCA1 mutant cells compared to intact junctions (white arrows) in hTERT control. N, nucleus. Scale bar = 2 μm.

In lesional skin, keratin fibers were reported to be aggregated around the nucleus in thick electron dense bundles that were also heavily stained by immunohistochemistry [18]. Perinuclear keratin (CK14) bundles were observed in SPCA1^-/-^ keratinocytes but were less apparent in SPCA1^-/+^ cells (**Fig. 2c**) although a few keratin bundles were visible on electron micrographs (**Fig. 2e**, ahead). Dysregulation of actin filaments has been widely reported in HHD, with acantholytic areas characterized by strong peripheral staining of F-actin arranged in parallel bundles beneath the cell membrane [18]. We observed similar peripheral staining with fluorescently labeled phalloidin in SPCA1 mutant cell lines together with some nucleated structures in the cytoplasm (**Fig. 2d**).

Electron microscopy of keratinocyte junctions revealed additional differences between hTERT (control) and SPCA1 mutant cells. In both low and high calcium medium, control keratinocytes showed smooth junctions with some microvilli making cell-cell contacts as described [36] (**Fig. 2e**). However, in both SPCA1^-/+^ and SPCA1^-/-^keratinocytes, there were pronounced intercellular gaps, some containing isolated cell fragments similar to the intercellular “bleb islands” observed by electron microscopy of cells from HHD lesions [18]. The intercellular voids and blebs failed to resolve in high Ca^2+^, consistent with acantholysis and defective cell–cell adhesion characteristic of HHD.

### SPCA1 is required for actin-mediated keratinocyte spreading

Previous studies reported abnormal actin organization, localization and stress fiber formation in skin specimen from HHD patients [18, 19] that persist in vitro [20]. However, actin-mediated changes in keratinocyte morphology were not explored. The actin cytoskeleton is required for maintenance and changes in cell shape, migration and spreading by mediating attachment to the extracellular matrix, formation of leading edge and dynamic remodeling of the retracting edge. Here, we assayed cell spreading by harvesting cells with trypsin and replating onto collagen coated plates in complete media or basal media lacking supplemental growth factors (**Fig. 3a**). Following replating, cells initially have round morphologies with small areas, but after cell attachment, membrane protrusions and new cell-matrix adhesions are formed that increase cell area resulting in cell spreading [37]. Four hours after replating, cells were fixed, permeabilized and stained with fluorescently labeled phalloidin to visualize filamentous actin (**Fig. 3b**). We observed significant differences in cell area between control keratinocytes and both complete SPCA1^-/-^ and heterozygous SPCA1^-/+^ knockouts (**Fig. 3c**). Interestingly, the cell spreading defect in SPCA1^-/+^ keratinocytes was similar (in complete media) or worse (in basal media) than that seen for SPCA1^-/-^cells. Furthermore, the difference in cell spreading in complete versus basal media observed in normal keratinocytes was not seen for SPCA1 mutant lines (**Fig. S3**).

**Figure 3.**
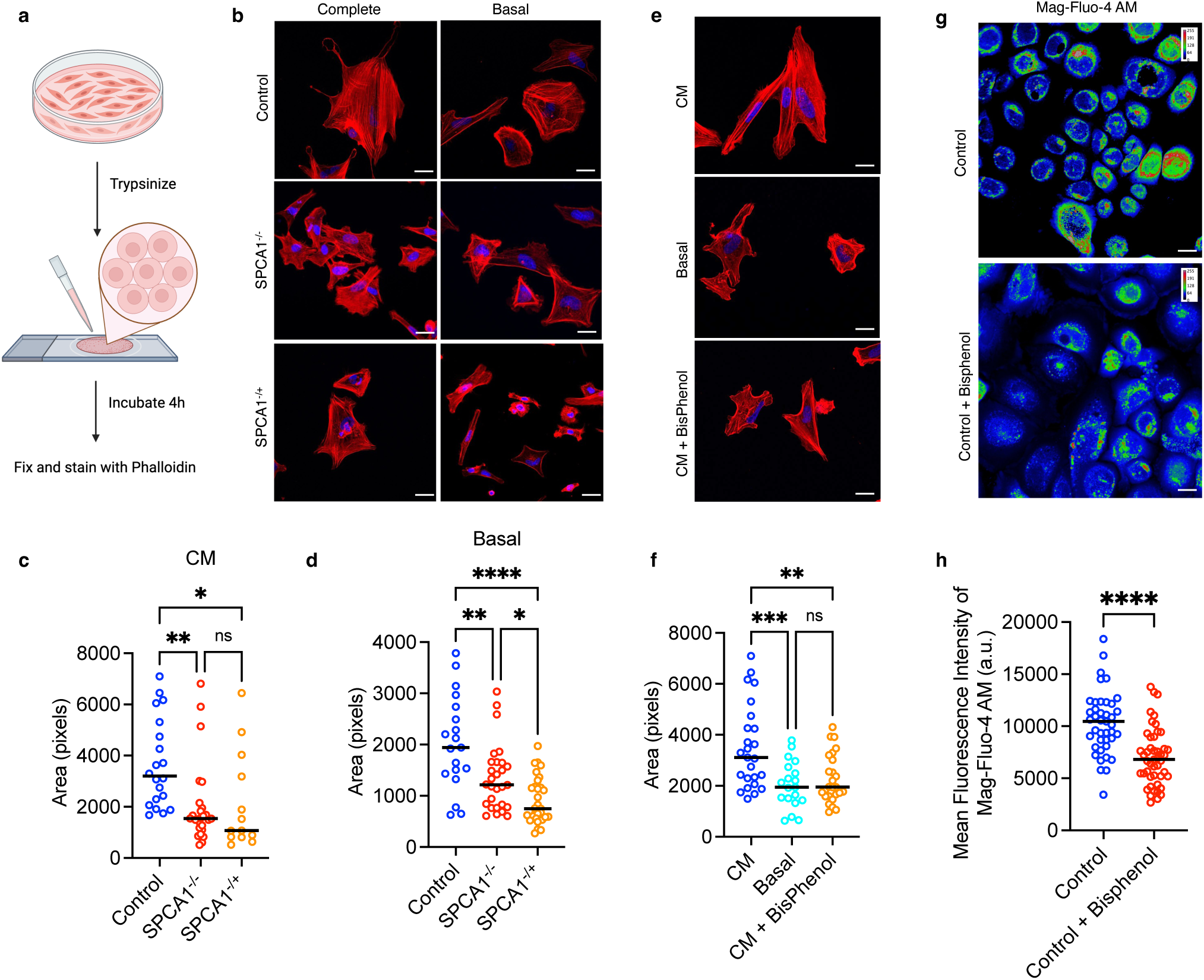
SPCA1 controls actin cytoskeletal dynamics required for keratinocyte spreading. **(a)** Schematic of the cell spreading assay. **(b)** Representative fluorescence images and **(c-d)** quantification of area as visualized by phalloidin staining after 4 h of recovery in either complete (CM) or basal media. Reduced membrane protrusions and impaired spreading was observed in SPCA1 mutants relative to hTERT keratinocytes. **(e)** Representative fluorescence images and **(f)** quantification of cell spreading in control keratinocytes treated with bisphenol (2.5 μM for 4 h). Keratinocyte spreading in complete medium (CM) after bisphenol treatment was not significantly different from basal media. Graphs represent mean ± SD for n = 25-35 cells per group. Data were evaluated using Ordinary One-way ANOVA analysis. ns = not significant; *\*P<0.05; **P<0.01; ***P<0.001; ****P<0.0001.* **(g)** Representative fluorescence images with **(h)** quantification of Mag-Fluo-4 AM in control keratinocytes in response to acute addition of bisphenol (5 μM). Bisphenol significantly reduced perinuclear Ca²⁺-sensitive fluorescence of Mag-Fluo-4 AM. Magnification, 40x. Scale bar = 20 μm. Graphs represent mean ± SD for n = 40-50 cells per group. Data were evaluated using unpaired t-test, *\*\*\*\*P<0.0001*.

To confirm the observed link between Golgi Ca^2+^-ATPase dysfunction and actin cytoskeleton reorganization, we evaluated the effect of the SPCA1-selective inhibitor bisphenol on cell spreading. Acute addition of bisphenol (5 μM) in MagFluo4-AM loaded normal keratinocytes resulted in significant reduction of Ca^2+^-sensitive fluorescence in the perinuclear region, as expected (**Fig. 3d-e**). Next, we evaluated the effect of bisphenol on cell spreading. In complete medium, bisphenol (2.5 μM for 24h) treated keratinocytes failed to regrow after trypsinization and plating on complete media (**Fig. 3f**), appearing similar in cell size to those plated on basal media (**Fig. 3g**). These observations, taken together with the ability of transgenic hSPCA1 to rescue cell spreading defects in the mutant keratinocytes (**Fig. 7**, ahead), point to a role for SPCA1 in actin cytoskeletal reorganization.

### Cell spreading in keratinocytes is driven by insulin

While essential nutrients such as amino acids, vitamins and trace minerals are supplied in basal media to support keratinocyte metabolism and growth, complete media additionally contains growth factors (EGF), hormones (insulin, hydrocortisone) and serum-containing extract (bovine pituitary extract). We noted (**Fig. S3**) that in normal keratinocytes, cell spreading was significantly enhanced in complete medium. First, we confirmed this observation using an alternative protocol to monitor dynamic changes in cell size. Keratinocyte cultures were placed in Hanks balanced salt solution (HBSS) for two hours, then changed to complete media for varying lengths of time (0 to 30 minutes) before being fixed and stained with phalloidin (**Fig. 4a**). As seen in the representative images (**Fig. 4b**) and quantifications (**Fig. 4c**), replacing complete media (“untreated”) with osmotically balanced HBSS resulted in cell shrinkage (“0 min”), but the keratinocytes quickly “regrow” within 30 min upon reintroduction of complete medium.

**Figure 4.**
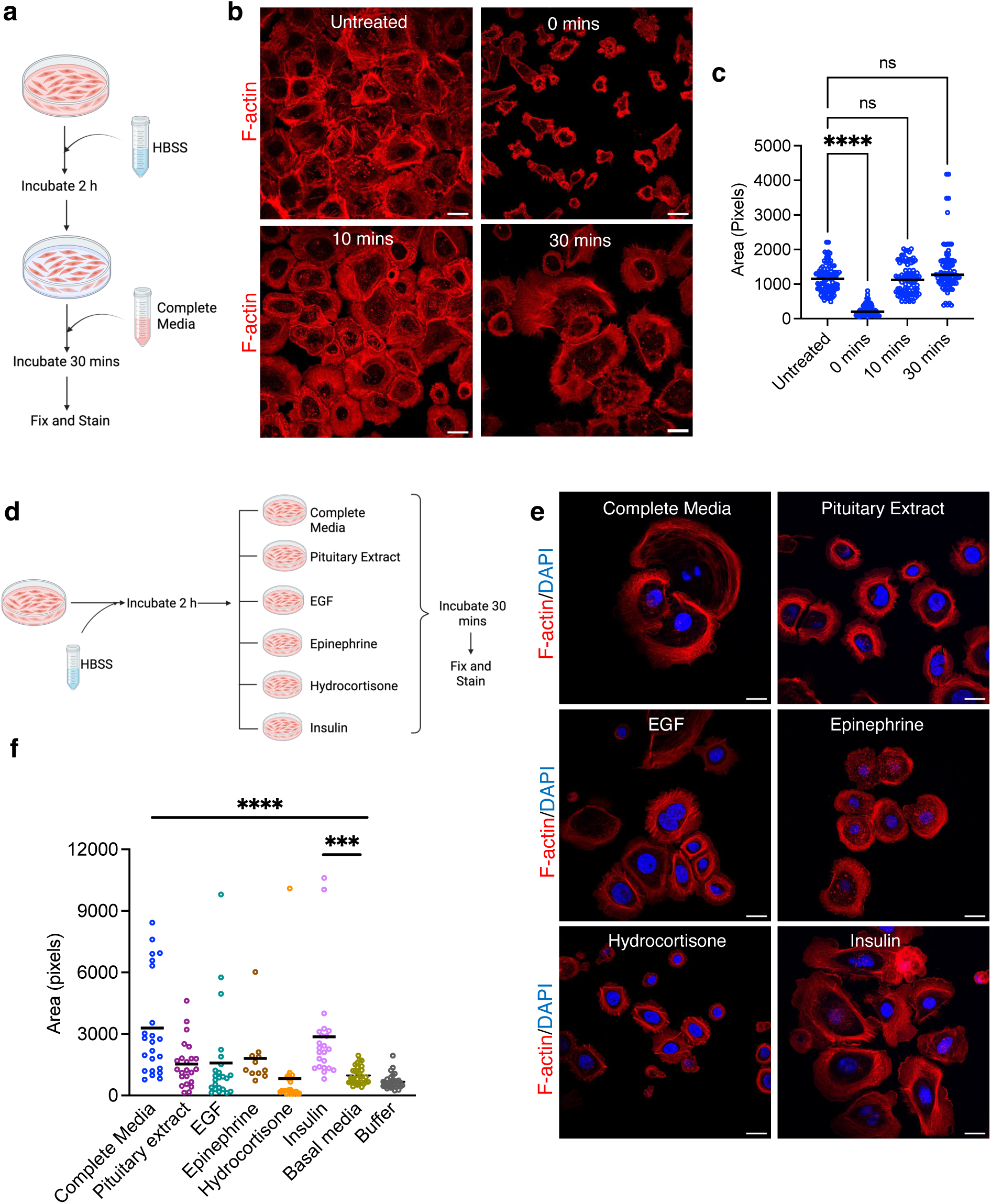
Keratinocyte spreading requires insulin signaling. **(a)** Schematic of the cell spreading assay. **(b)** Representative images and **(c)** quantification show keratinocyte shrinkage in HBSS and rapid recovery of cell area following reintroduction of complete medium. Magnification, 20x. Scale bar = 20 μm. Graph represent mean ± SD for n = 90 cells per group. **(d)** Schematic of cell spreading assay to determine the effect of individual components. **(e)** Representative images and **(f)** quantification of cell area on addition of individual supplemental components to basal media. Insulin alone restored cell spreading to levels comparable to complete medium. Differences in cell spread between other individually supplemented media and basal media were not significant. Magnification, 20x. Scale bar = 20 μm. Graph represent mean ± SD for n = 11-30 cells per group. Data were evaluated using Ordinary One-way ANOVA analysis. *\*\*\*P<0.001; ****P<0.0001*.

Next, we added supplemental factors individually to basal media, as shown (**Fig. 4d**). Addition of insulin alone to basal medium was sufficient to support cell spreading as seen in complete medium, whereas other supplemental additions to basal media did not significantly increase cell size when compared to basal medium or HBSS buffer alone (**Fig. 4e-f**). The dependence on insulin is consistent with previous data that showed a role for insulin like growth factor (IGF), but not epidermal growth factor (EGF), in membrane protrusion and keratinocyte spreading [38].

### SPCA1 is required for insulin-mediated cell spreading and lamellipodia formation

We asked if the cell spreading defect in SPCA1^-/-^ and SPCA1^-/+^ keratinocytes observed in complete medium (**Fig. S3**) could be attributed to insulin-mediated processes. Keratinocytes were trypsinized and plated on collagen-coated basal medium supplemented only with insulin for 4 h as described in **Fig. 3a**. Control keratinocytes stained with phalloidin showed long spike-like extensions that were notably absent in both SPCA1^-/-^ and SPCA1^-/+^ keratinocytes, suggesting a defect in insulin response (**Fig. 5a**). Next, keratinocyte cultures were placed in HBSS for two hours, then changed to basal medium supplemented only with insulin for 30 min according to the schematic shown in **Fig 4d**. Control keratinocytes displayed prominent lamellipodia, which are sheet like projections that propel the leading edge of the cell (**Fig. 5b**, *box*). Lamellipodia were conspicuously absent or smaller in SPCA1^-/-^ and SPCA1^-/+^ keratinocytes, respectively. On the other hand, thin actin-based protrusions beyond the leading edge known as filopodia were observed in all samples and appeared more prominent in the SPCA1 mutants (**Fig. 5b**). For live imaging of actin dynamics, we transfected keratinocytes with Lifeact, a short actin binding peptide that is fused with GFP [39]. As seen in the supplemental movies (**Fig. S4a-c**) represented in the still image shown in **Fig. 5c**, the ebb and flow of dynamic cortical actin was evident in control keratinocytes but nearly absent or greatly diminished in SPCA1^-/-^ and SPCA1^-/+^ keratinocytes, respectively.

**Figure 5.**
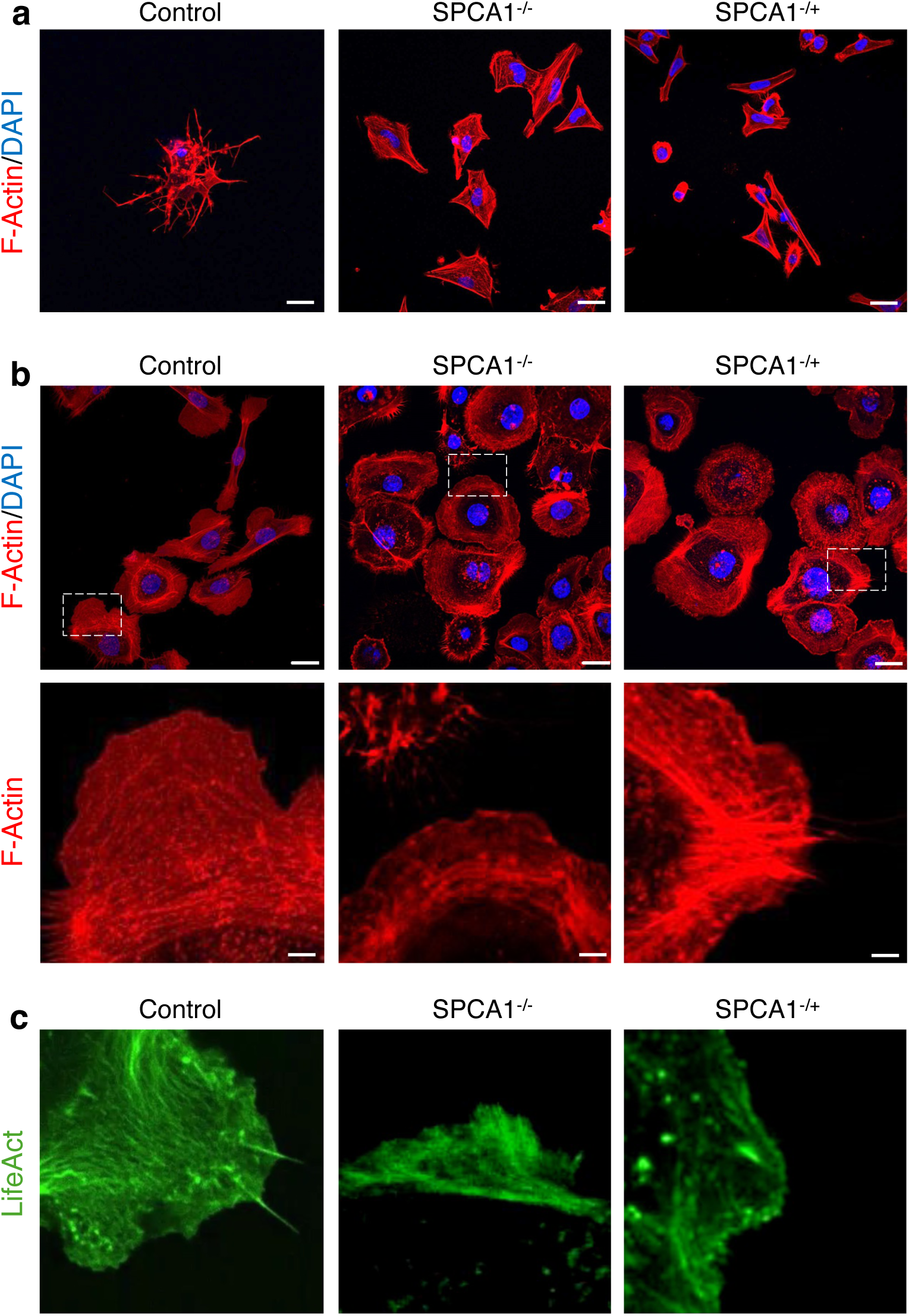
Insulin-induced keratinocyte spreading and lamellipodia formation requires SPCA1. Representative fluorescence images of phalloidin stained F-actin in keratinocytes **(a)** after being re-plated on collagen-coated plates for 4 h in basal media supplemented with only insulin. Control cells developed elongated, spike-like protrusions, absent in SPCA1 mutants, indicating impaired response to insulin. Magnification, 40x. Scale bar = 20 μm; **(b)** Similar to **(a)**, after 2 h incubation in HBSS and a 30 min recovery in basal medium supplemented only with insulin. Control cells displayed prominent lamellipodia at the leading edge which were absent in SPCA1^-/-^cells or reduced in SPCA1^+/-^ cells. Filopodia were observed in all genotypes and appeared more pronounced in SPCA1 mutants. Magnification, 40x. Scale bar = 10 μm; **(c)** Representative frame taken from live cell imaging shown in **Fig. S4** of Lifeact-GFP–expressing keratinocytes showing well-formed lamellipodia in control cells but not in SPCA1 mutants. See **Fig. S4** for differences in actin dynamics between the three keratinocyte lines.

### Insulin signaling pathway leading to Rac1 GTPase activation is defective in SPCA1 mutant keratinocytes

The insulin signaling pathway is activated by insulin binding to its membrane receptor and initiating a cascade of events that lead to actin cytoskeleton reorganization and membrane ruffling, via phosphorylation of the PI3K and AKT kinases and activation of the small GTPase Rac1 [40–42] (**Fig. 6a**). We investigated key steps involved in insulin signaling in normal and SPCA1 mutant keratinocytes. Expression levels of the insulin receptor were significantly lower than control in the SPCA1^-/-^ null mutant but appeared normal in heterozygous SPCA1^-/+^ keratinocytes (**Fig. 6b**). However, levels of the downstream phospho-AKT protein were similarly depressed in both SPCA1 mutant lines (**Fig 6c**). We used FITC-labeled insulin to monitor uptake into cells (**Fig. 6d**). Consistent with observed differences in levels of insulin receptor, uptake of FITC-insulin was more significantly decreased in the SPCA1 null keratinocytes, compared to the heterozygous mutant. PI3K is typically upstream of AKT kinase activation. Therefore, to determine a potential role for PI3K in cell spreading, we treated keratinocytes with LY294002, a PI3K inhibitor used at a concentration that does not affect other kinases [40]. Insulin-stimulated keratinocyte spreading was completely inhibited by LY294002 (**Fig. 6e**).

**Figure 6.**
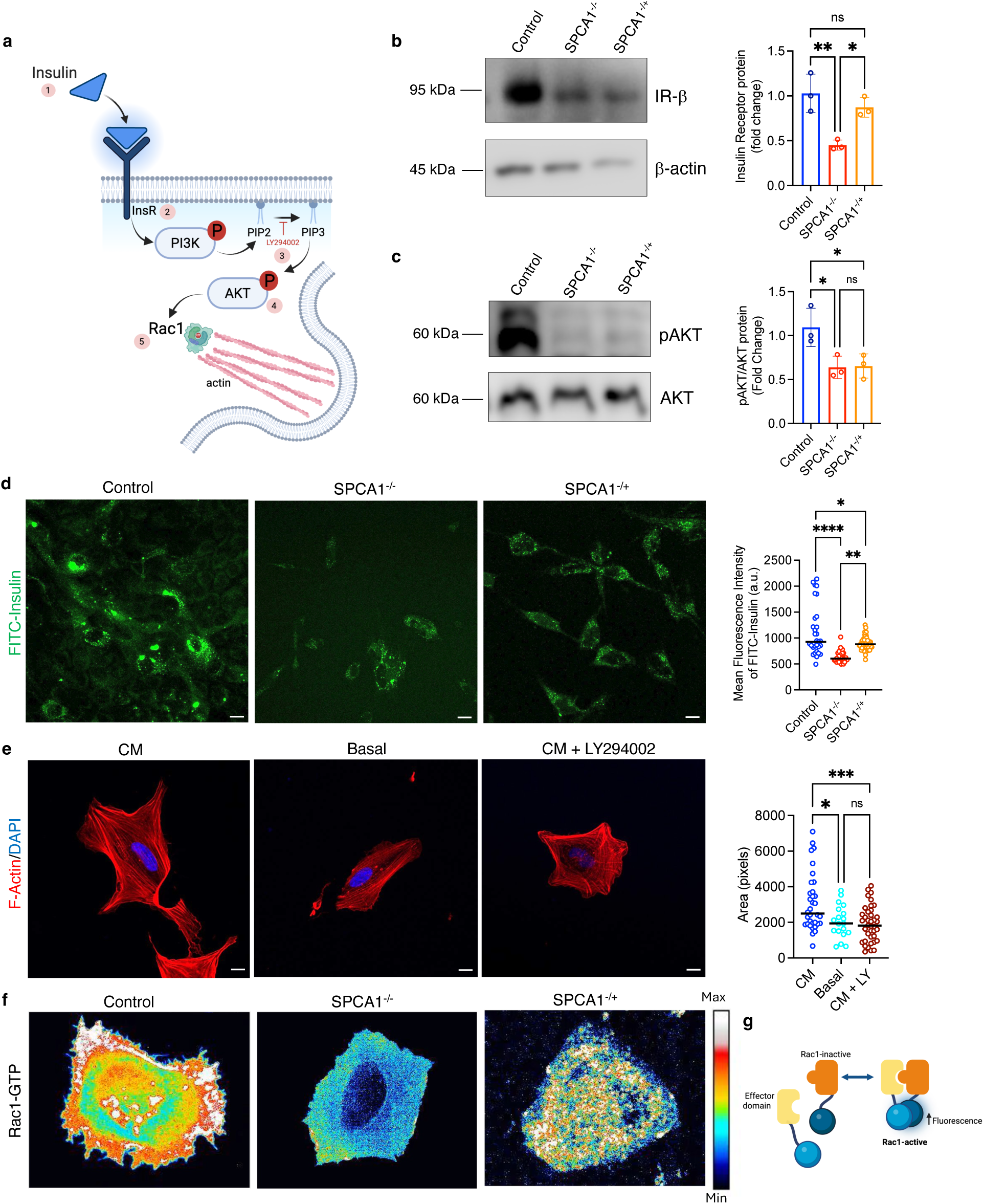
SPCA1 loss impairs PI3K–AKT signaling and Rac1-Mediated actin remodeling. **(a)** Schematic of the insulin signaling pathway leading to PI3K/AKT activation and Rac1-driven actin remodeling. Numbers show the order of events. **(b-c)** Representative Western blot images and quantification of **(b)** Insulin receptor (IR-β) normalized to β-actin as loading control and **(c)** pAKT levels relative to total AKT. Western blot analysis shows reduced insulin receptor levels in SPCA1^-/-^ keratinocytes, while SPCA1^+/-^ cells retain near-normal expression. Phosphorylation of AKT is markedly decreased in both SPCA1 mutant lines. Graphs represent mean ± SD for n=3 replicates per group. **(d)** Fluorescence images of internalized FITC-insulin in keratinocytes. Uptake was significantly diminished in SPCA1^-/-^ keratinocytes and moderately reduced in SPCA1^+/-^ cells, consistent with differences in receptor levels. **(e)** Representative fluorescence images of phalloidin stained F-actin and quantification of keratinocyte area of control keratinocytes in complete media treated with PI3K inhibitor (50 µM LY294002 for 1 h) before stimulating with insulin. LY294002 (LY) completely blocked insulin-stimulated keratinocyte spreading. Magnification, 20x. Scale bar = 10 μm. Graphs represent mean ± SD for n = 20-60 cells per group. **(f)** Representative still images of Rac1 GTPase activation taken from the live cell movies (**Fig. S5**) visualized by a dimerization-dependent fluorescent biosensor schematically depicted in **(g)**. Insulin robustly activated Rac1 in control keratinocytes but elicited weak or absent activation in SPCA1 mutants. Data were evaluated using Ordinary One-way ANOVA analysis. ns = not significant; *\*P<0.05; ***P<0.001; ****P<0.0001*.

The RhoA family of small GTPases, including RhoA, Cdc42 and Rac1, play critical roles in cell motility. Previous studies have shown that RhoA is not involved in insulin-stimulated keratinocyte migration [40]. Rac1 is an important target of PI3K and a major control point for actin cytoskeleton reorganization and lamellipodia formation [42, 43]. To dynamically monitor Rac1 activation with high spatiotemporal resolution we used a dimerization dependent fluorescent biosensor wherein each quenched monomer is attached to either the small GTPase or effector domain [44] (**Fig. 6f**, *right*). In response to insulin, activity dependent interaction of the Rac1 GTPase and its effector results in fluorescence as captured in the still images shown in **Fig. 6f** and in the movies shown in **Fig. S5a-c**. Fluorescence intensity corresponding to Rac1 activation was sharply decreased in the SPCA1 mutants, compared to the robust response in control cells. In summary, key steps in the insulin signaling pathway were found to be defective in the SPCA1 mutants, with a graded response from the single to double allele knockouts.

### Transgenic hSPCA1 or Ca^2+^-ATPase agonist reverse HHD cellular phenotypes

As proof of concept for future gene therapy approaches, we transfected hSPCA1 into SPCA1^-/+^ keratinocytes and assessed reversal of key pathological phenotypes. As expected, we observed increased hSPCA1 expression colocalizing with GM130 in Golgi stacks that appeared more compact in the transfected cells (**Fig. 7a**), consistent with the rescue of Golgi fragmentation phenotype. Western blotting revealed only modest increase in SPCA1 expression in SPCA1^-/+^ keratinocytes (**Fig. 7b-c**), likely due to the limited accessibility of keratinocytes to transient transfection. Despite this, functional rescue of the cell spreading defect in SPCA1^-/+^ keratinocytes was evident. Following trypsinization and replating on collagen, transfected cells appeared similar to normal hTERT keratinocytes and showed more significant regrowth in both complete and basal media compared to the SPCA1^-/+^ keratinocytes (**Fig. 7d-f**). Transfected cells also recovered the ability to uptake FITC-insulin, appearing indistinguishable to hTERT keratinocytes (**Fig. 7g-h**).

**Figure 7.**
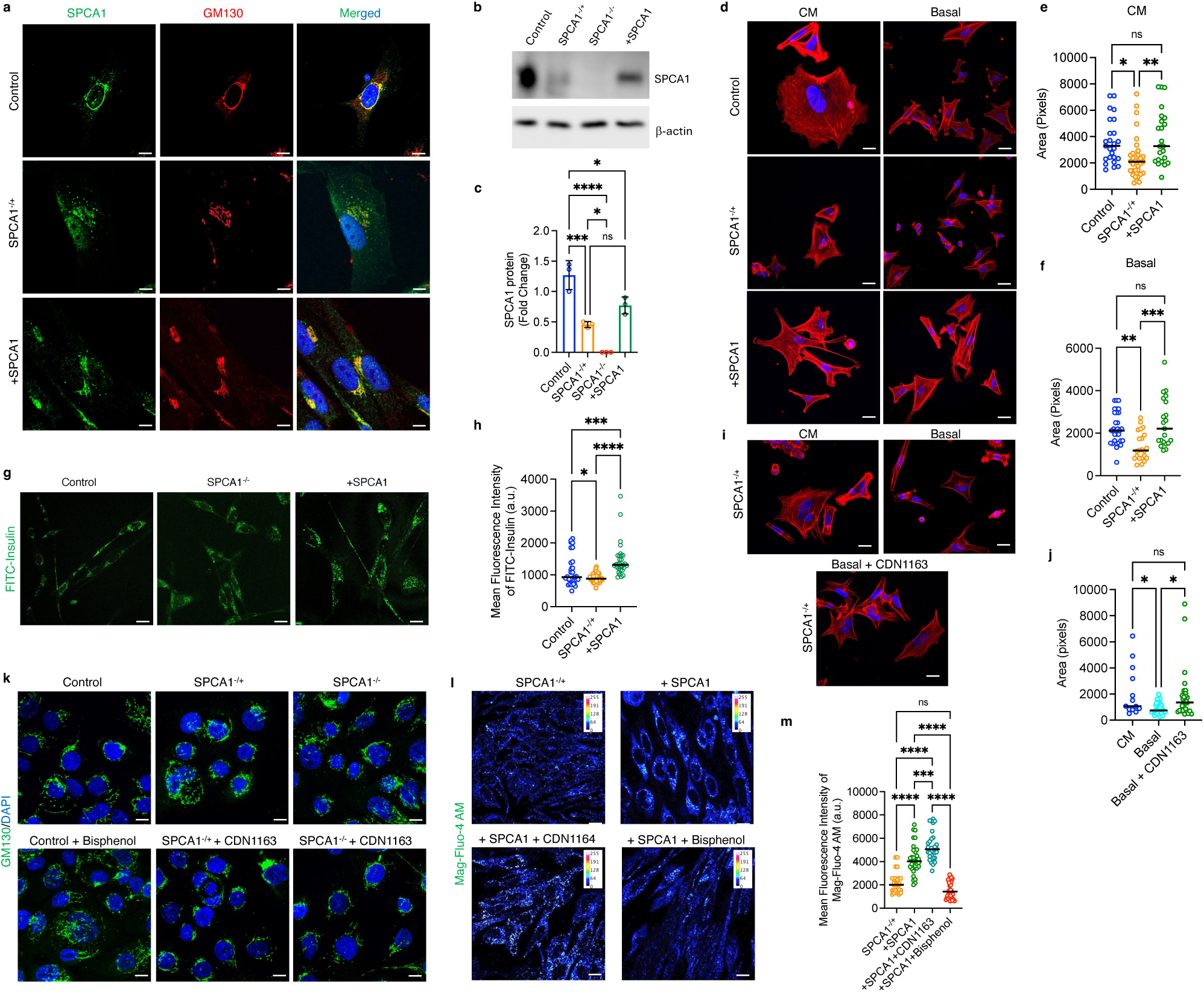
Transgenic expression or activation of Ca²⁺-ATPase rescues HHD phenotypes. **(a)** Representative immunofluorescence images of GM130 and SPCA1 in keratinocytes. Magnification, 63x. Scale bar = 5 μm. SPCA1^+/-^ keratinocytes transfected with hSPCA1 exhibit increased SPCA1 expression and more compact Golgi morphology. **(b)** Representative Western blot and **(c)** quantification of total levels of SPCA1 relative to β-actin loading control. **(d-f)** Representative fluorescence images **(d)** and **(e-f)** quantification of cell area visualized by phalloidin staining after 4 h of recovery in either complete or basal media as described in Fig. 3A. Functional rescue of the cell-spreading defect is observed in hSPCA1-transfected SPCA1^+/-^ keratinocytes, which gain the ability to regrow, after replating, to levels comparable to hTERT control. Magnification, 40x. Scale bar = 10 μm. Graphs represent mean ± SD for n = 20-60 cells per group. **(g)** Representative images and **(h)** quantification of FITC-insulin uptake in keratinocytes. hSPCA1 transfection in SPCA1^+/-^ cells restored FITC-insulin uptake, similar to control. Magnification, 40x. Scale bar = 10 μm. Graphs represent mean ± SD for n = 30-60 cells per group. **(i)** Representative fluorescence images and **(j)** quantification of cell spreading capacity of control keratinocytes in presence of Ca^2+^-ATPase agonist CDN1163 (2.5 μM for 4 h) as indicated. Cell spreading in basal medium after CDN1163 treatment was similar to that in complete medium. **(k)** Golgi morphology, seen by GM130 immunofluorescence in keratinocytes. Golgi fragmentation in hTERT cells was induced using bisphenol. Golgi fragmentation in SPCA1 mutant keratinocytes is largely reversed by CDN1163. **(l)** Representative fluorescence images and **(m)** quantification of Mag-Fluo-4 AM fluorescence in SPCA1^+/-^cells; hSPCA1-transfected SPCA1^+/-^ cells (+SPCA1); hSPCA1-transfected SPCA1^+/-^ cells acutely treated with 5 µM CDN1163 (+SPCA1+CDN1163) or 5 µM Bisphenol (+SPCA1+Bisphenol) for 30 min. Magnification, 40x. Scale bar = 20 μm. Graph represents mean ± SD for n = 30-35 cells per group. Data were evaluated using Ordinary One-way ANOVA analysis. ns = not significant; *\*P<0.05; ***P<0.001; ****P<0.0001*.

These results prompted us to evaluate a small molecule Ca^2+^-ATPase agonist, CDN1163, that has been reported to enhance the activity of the related sarco/endoplasmic reticulum Ca^2+^-ATPase (SERCA) [45, 46]. SPCA1^-/+^ keratinocytes in basal medium supplemented with CDN1163 showed actin cytoskeleton-mediated regrowth similar to that in complete medium (**Fig. 7i-j**). Golgi morphology is known to be regulated by Ca^2+^ homeostasis [47]. We show that treatment of hTERT keratinocytes with the SPCA1 inhibitor bisphenol results in fragmentation of Golgi stacks, as revealed by confocal microscopy (**Fig. 7k**) and 3D reconstruction (**Fig. S1**) of GM130 immunofluorescence. Golgi fragmentation in SPCA1 mutants was largely reversed by CDN1163, in both single and biallelic *ATP2C1* knockouts (**Fig. 7k** and **Fig. S1**). These findings suggest that CDN1163 boosts Ca^2+^ homeostasis in SPCA1 mutant cells. To directly gauge the effect of CDN1163 on Golgi Ca^2+^ stores, we evaluated MagFluo-4 AM fluorescence in these cells (**Fig. 7l-m**). In SPCA1-/+ keratinocytes, MagFluo-4 AM fluorescence was sparse and scattered, consistent with fragmented Golgi and depleted Ca^2+^ stores described earlier in **Fig. 1**. Ectopic expression of hSPCA1 not only increased fluorescence but also consolidated the fluorescence in crescent shaped structures reminiscent of the Golgi organelle. Ca^2+^-dependent fluorescence was further enhanced in these cells by CDN1163 albeit with a more dispersed pattern, consistent with the endoplasmic reticulum. Finally, treatment with bisphenol diminished MagFluo-4 fluorescence, reversing the effect of transgenic SPCA1. These observations confirm and extend the applicability of calcium pump activation in ameliorating a range of disorders. In summary, we conclude that gene therapy and/or small molecule agonists may be promising avenues for therapeutic resolution of HHD.

## Discussion

The HHD phenotype is consistent with SPCA1 haploinsufficiency, with more than 50% of 82 HHD mutations reported in one study associated with lower levels of *ATP2C1* transcript [12]. Skin samples and keratinocytes obtained from HHD patients showed reduced levels of SPCA1 expression, described as weak to moderate staining by immunohistochemistry [48, 49], and averaging 50% reduction by Western blotting [17]. Based on these observations, the single allele CRISPR/Cas9-mediated *ATP2C1* disruption described here (SPCA1^-/+^), showing 50% reduction in SPCA1 protein expression could be a good model for HHD. Whereas typical HHD results from germline mutations and exhibits a generalized, bilateral involvement, type 1 segmental HHD reflects a localized post-zygotic mutation in the early stages of embryogenesis, conferring segmental cutaneous phenotypes. A further (type 2) segmental form results from a combination of both the germline mutation and a somatic loss of heterozygosity in the embryo, resulting in lesions with both hemizygous and homozygous *ATP2C1* mutations [50]. The loss of heterozygosity leads to more severe HHD phenotypes appearing earlier in life [51]. It was thus useful to evaluate the biallelic *ATP2C1* knockout (SPCA1^-/-^) as well.

A comparison of heterozygous and homozygous *ATP2C1* null mutants led to some intriguing observations. While SPCA1^-/+^ keratinocytes displayed intermediate phenotypes in SPCA1 expression, Golgi morphology and Ca^2+^ accumulation as expected, downstream effects of the mono-allelic *ATP2C1* disruption sometimes resembled the bi-allelic knockouts or even appeared more severe. This was apparent in the near complete loss of cadherins in the desmosomes and adherens junctions in both SPCA1^-/+^ and SPCA1^-/-^ keratinocytes. Functional dysregulation of actin cytoskeleton mediated cell spreading and insulin signaling were varied, with the heterozygous mutant exhibiting some intermediate phenotypes and others similar to the full knockout. These observations hint at complex multi-pathway interactions that underly the wide spectrum of HHD defects [52].

We show that loss of SPCA1 attenuates keratinocyte spreading, a critical step in wound healing. The process of reepithelialization begins with cells at the wound edge extending actin filament-powered lamellipodia to initiate migration and is driven by signaling molecules and growth factors secreted by immune cells and the surrounding tissue. Cadherin based interactions with adjoining cells and extracellular matrix ensure coordinated and efficient movement. Each of these critical steps is compromised in the SPCA1 mutants, beginning with defective cell junctions, growth factor driven signaling, actin cytoskeleton reorganization, Rac1 GTPase activation and lamellipodia formation. These findings not only exemplify the central importance of Ca^2+^ homeostasis in multi-pathway regulation but also reveal the critical role of SPCA1 in mediating these pathways.

The efficacy of insulin both topically and systemically in promoting wound healing has been known for nearly a century [53] although its mechanistic basis is still emerging. Insulin is a peptide hormone widely known for its role in regulating blood glucose levels. Although popular for its non-diabetic use in the early part of the 20^th^ century, therapeutic applications of insulin were ignored through the 40’s and 50’s but have revived in more recent years [40]. Insulin injections or topical applications have been demonstrated to improve healing of bone, burns, incision wounds and cutaneous ulceration both in experimental settings and in the clinic. Diabetic patients have been successfully treated with insulin spray. However, insights into the mechanism underlying the therapeutic use of insulin in wound healing are sparse. Using a model of excision wounds in mouse and invitro keratinocyte scratch assay, Liu et al. demonstrated the effect of insulin via the insulin receptor, but not the EGF-receptor, through the PI3K-AKT-Rac1 pathway [40]. Together with this work emphasizing the role of Ca^2+^ signaling and the Golgi/secretory pathway Ca^2+^-ATPase in insulin-mediated keratinocyte spreading, the case for insulin as a powerful therapy for regenerative wound repair and restoration of the barrier function of skin is greatly strengthened. The potential application of insulin in these settings is boosted by its long history of safe use and low cost. In contrast, other growth factors like TGF-β and EGF are prohibitively expensive to produce and have therefore not been successfully harnessed for therapeutic use.

In conclusion, SPCA1 emerges as a target not only for treatment of HHD, a monogenic and rare disorder, but also for more widespread application in treatment of a range of wounds in diabetic and non-diabetic patients. Our observations could spur the discovery of small molecule activators of SPCA1 that could be used in selectively or in combination with SERCA2 agonists. Further, we highlight the potential use of gene therapy approaches that could boost SPCA1 expression and activity in keratinocytes.

## Supporting information

All Supplemental Figures

## Ethics Statement

Not applicable.

## Data Availability Statement

All primary data will be made available upon reasonable request.

## Acknowledgments

The authors thank Dr. Luis Garza for helpful advice. This manuscript is the result of funding in whole or in part by the National Institutes of Health (NIH). It is subject to the NIH Public Access Policy. Through acceptance of this federal funding, NIH has been given a right to make this manuscript publicly available in PubMed Central upon the Official Date of Publication, as defined by NIH. Research reported in this work was supported by the following awards: National Center for Advancing Translational Sciences of the National Institutes of Health under award number 1R03TR005427 to RR and National Institutes of Arthritis and Musculoskeletal and Skin Diseases under award number 1P30AR075043 to JEG. Portions of the study were also supported by Formation Venture Engineering Foundry to RR. The content is solely the responsibility of the authors and does not necessarily represent the official views of the National Institutes of Health.

## Author Contributions Statement (CRediT-compliant)

Conceptualization: RG, AC, PNP, MKS, JEG, AV and RR; Data Curation: RG, AC, PNP, MKS; Formal Analysis: RG, AC, PNP, MKS; Funding Acquisition: RR, JEG, AV; Investigation: RG, AC, PNP, MKS, JEG, AV and RR; Methodology: RG, AC, PNP, MKS; Project Administration: RR and JEG; Resources: RR and JEG; Supervision: RR, AV and JEG; Validation: RG, AC, MKS and PNP; Visualization: RG, AC, PNP; Writing – Original Draft Preparation: RR; Writing – Review and Editing: RR, JEG, MKS, RG, AC and PNP.

## Materials and Methods

### Cell lines and culture

*ATP2C1* gene knockouts were engineered in human epidermal keratinocytes from original line N/TERT-2G [54]. CRISPR KO KCs were generated as previously described [55]. In brief, single-guide RNA (sgRNA) target sequence was developed (*ATP2C1* sgRNA: GGAGCTGTCACCTTAGAACA for *ATP2C1* KO using a CRISPick (https://portals.broadinstitute.org/gppx/crispick/public), web interface for CRISPR sgRNA. Synthetic sgRNA target sequences were inserted into a cloning backbone, pSpCas9 (BB)-2A-GFP (PX458) (Addgene # 48138), and then cloned into One Shot Stbl3 chemically competent *E. coli* cells (Thermo Fisher Scientific # C737303). Proper insertion was validated by Sanger sequencing. The plasmid with proper insertion was then transfected into an immortalized KC line (N/TERT-2G) using the TransfeX transfection kit (ATCC # ACS4005) in the presence of JAK1/JAK2 inhibitor, baricitinib (10 μg/ml, AchemBlock # G-5743). GFP-positive single cells were plated and then expanded. Cells were then genotyped and analyzed by Sanger sequencing using these primers, ATP2C1PCRF1: TGTTCCCTAAGGCTGTGGATATP2C1PCRR1: CTTTGCTTTGCCACATCTGA. Homozygous and heterozygous *ATP2C1* KO keratinocytes were validated by Western blot (ATP2C1 monoclonal antibody, Abnova # H00027032).

Cells were grown in KGM^®^ Gold Keratinocyte Growth Medium BulletKit (Lonza #0019260) supplemented with 10% heat inactivated FBS and 100X antibiotic-antimycotic solution (Thermo Fisher Scientific). The cells were grown at 37°C in a humidified atmosphere containing 5% CO_2_. For all the experiments, cells were grown on collagen coated tissue culture plates to promote cell adherence.

### DNA transfection

SPCA1^-/+^ cells were transfected with 1 µg hSPCA1 overexpression plasmid DNA using Polybrene method. Briefly, hSPCA1 plasmid DNA was incubated in antibiotics free KBM complete medium containing 4 µg/mL Polybrene transfection reagent. The mixture was incubated at RT for 5 mins and then added to cells for 24 h at 37°C. After incubation, cells were left to recover for 24 h in antibiotics free KBM complete medium at 37°C. Post-recovery, cells were harvested and seeded for experiments as required.

### Protein extraction and Western blotting

Proteins were extracted from cultured cells by incubating cells in RIPA buffer (Thermo Fisher Scientific #89901) supplemented with 100x protease/phosphatase inhibitor cocktail (Cell Signaling #5872S) on ice for 30 mins. After incubation, supernatants were collected after centrifugation at 13000×g for 5 minutes. Supernatant was collected in fresh 1.5 mL Eppendorf tubes. Protein quantification was done using Pierce BCA Protein Assay Kit (Thermo Scientific # 23227). For each sample, 20 µg of total protein was resolved by SDS-Polyacrylamide gel electrophoresis using 8% Bis-Tris gels. The separated proteins were then transferred to PVDF membrane and blocked in 1x TBST containing 2% BSA for 1 h at room temperature (RT). Immunoblots were incubated overnight at 4°C with primary antibodies from Cell Signaling Technologies: pAKT (#4060P); AKT (#9272S); Insulin receptor-β (23413S); *β*-actin (4967S); from Abnova: ATP2C1 (H00027032); and from Invitrogen: E-cad (#13-1700), followed by incubation with HRP-conjugated secondary antibodies for 1 h at RT. Signal was detected using chemiluminescence reagent Immobilon^®^ Forte Western HRP-Substrate (Millipore #WBLUF0500) and the blots were analyzed using ImageJ. Alternatively, IRDdye secondary goat antibodies (Li-Cor Biosciences), anti-mouse (680RD, Cat. no. 926–68070), and anti-rabbit (800CW, Cat. no. 926–32211) were used at a 1:20,000 dilution for 1 hour at RT. The blots were visualized and quantified with the Odyssey System (Li-Cor Biosciences).

### Immunofluorescence

#### Intracellular labeling

Cells grown on coverslips were fixed with 4% paraformaldehyde (PFA) for 10 mins at RT and then washed thrice with PBS. Cells were then permeabilized with 0.25% Triton X-100 for 15 mins at RT, followed by 1 h blocking with 1x PBS containing 5% BSA and 2% normal goat serum. Primary antibodies against ATP2C1, GM130, Cytokeratin 14 (CK-14) (Proteintech #60320) and E-cad were incubated overnight at 4°C or at least 1 h at RT in the blocking buffer at 1:50, 1:300, 1:100 and 1:100 dilution, respectively. After incubation, cells were washed with PBS and incubated with Alexa 488- or Alexa 568-conjugated secondary antibodies diluted at 1:2000 for 1 h at RT. Cell nuclei were stained blue using DAPI. Coverslips were mounted on glass slides using prolong Gold antifade reagent (Invitrogen #P36935). Immunostaining was visualized using an inverted Zeiss LSM700 confocal microscope.

#### Cytoskeleton labeling

Cells grown on coverslips were fixed with 4% PFA for 10 mins at room temperature followed by permeabilization with 0.25% Triton X-100 for 5 mins at RT. Cells were washed with PBS followed by incubation with 180nM solution of Rhodamine phalloidin-TRITC (Hello Bio #HB8621) in 1x PBS containing 1% BSA for 1 h at RT. Cells were washed with PBS, nuclei were stained blue using DAPI and mounted on glass coverslips for visualization using Zeiss LSM700 confocal microscope.

To study the effect of Ca^2+^ on cell junctions, after cells reached confluency, cells were incubated with complete medium containing 1.2 mM Ca^2+^ for 24 h. Cells were fixed with 4% PFA post-incubation and stained with TRITC-phalloidin as described above.

### Time lapse imaging

Cells were transfected with RHPMP192 (pLifeAct sfGFP) plasmid (Addgene #88843) using Lipofectamine 3000 as per manufacturer’s instructions (Thermo Fisher Scientific). Cells expressing the plasmid were grown on glass bottom 35 mm petri-dishes and imaged using Nikon SoRa confocal microscope in lattice structured illumination microscopy (SIM) mode at 5 s interval for 2 min 25 s. Kymographs of these images were generated using the inbuilt Fiji plugin ‘Multi Kymograph’ and selecting line width as ‘5’.

### MagFluo4-AM Fluorescence

Cells cultured in 35 mm glass-bottom collagen-coated dishes (MatTek # P35GCOL-1.5-14-C) to ∼80% confluency were washed in calcium and magnesium-free HBSS (Gibco #14170112). The cells were loaded with Mag-Fluo-4 AM (AAT Bioquest #20401) in calcium and magnesium-free HBSS at a final concentration of 3 µM with 0.02% Pluronic® F-127 (Invitrogen #P3000MP) at 37°C for 60-90 mins. After incubation, the cells were washed and resuspended in calcium-free imaging buffer (125 mM NaCl, 5 mM KCl, 1 mM MgCl2, 5 mM glucose, 25 mM HEPES, 0.1% w/v BSA, pH 7.4) containing 0.5 mM probenecid (Thermo Fisher Scientific # P36400). The cells were visualized using an inverted Zeiss LSM700 confocal microscope, and the images were analyzed using ImageJ.

### Electron microscopy

Cells were fixed with 2.5% glutaraldehyde in 3 mM MgCl_2_, 0.1M sodium cacodylate buffer. After rinses in buffer, cells were post-fixed for 1 h in 1% osmium tetroxide. Samples were rinsed in water then dehydrated in a graded series of ethanol followed by Embed-812 epoxy resin. The next day, resin was changed out for fresh resin every 2 h, and then the plate was dried in an oven set at 60°C. After the resin (containing cells) was solidified, it was removed from the plate and trimmed to blocks for sectioning. Ultrathin (60-70 nm) sections were cut using a Leica UC7 ultramicrotome and collected on copper grids, followed by staining with UranyLess and Lead Citrate. TEM reagents were purchased from Electron Microscopy Sciences. TEM imaging was performed using a ThermoFisher Talos L120C at 120KV and acquired with a Ceta CMOS camera.

### Cell Spreading and actin regrowth assay

Cells were harvested and seeded at a density of 1.5 x 10^5^ cells per well on gelatin-coated coverslips, followed by incubation in either complete medium or basal medium, with or without LY294002 (PI3K inhibitor), Bisphenol (SPCA1 inhibitor), or CDN1163 (SERCA activator), for 4 h at 37°C. After incubation, cells were fixed with 4% PFA followed by cytoskeleton labelling with Rhodamine phalloidin-TRITC as explained earlier. As an alternative to the method described above, cells were grown to 80% confluency on coverslips. On the day of the experiment, cells were shrunk by replacing the cell media with nominally Ca free isotonic HBSS buffer for 2 h at 37°C. After incubation, complete media was added back for a period of 0-30 mins to investigate the rate of actin regrowth. At designated time intervals, cells were fixed with 4% PFA followed by cytoskeleton labelling with Rhodamine phalloidin-TRITC. The role of individual components on actin regrowth was also investigated by incubating cells with basal medium containing only respective individual components for 30 mins at 37°C after cell shrinkage with nominally Ca free isotonic HBSS buffer as described above.

### Insulin uptake assay

Cells were grown to 80% confluency in 35mm glass bottom petri-dishes. On the day of the experiment, cells were serum starved in basal medium for 2 h at 37°C. After incubation, 100 nM FITC-Insulin (Nanocs #IS1-FC-1) solution was added to cells for 15 mins at 37°C. Cells were then gently washed twice with PBS and imaged immediately using Zeiss LSM700 confocal microscope.

### Rac1 GTPase Activation

Glass-bottom dishes were coated with 100 µg/mL collagen for 1 hour at 37 °C before cell seeding. After washing the dishes with PBS, 1 × 10⁵ keratinocytes were plated. Cells were allowed to attach for 24 hours and then transfected with the pCMV-G-Rac1 plasmid (Addgene #180575) using Lipofectamine 3000, according to manufacturer’s instructions (Thermo Fisher Scientific). Following an additional 24-hour incubation, cells were imaged using the structured illumination microscopy (SIM) mode of Nikon SoRa microscope at 5-second intervals for a total duration of 2 min 25 s.

### Statistical Analysis

Statistical analyses were performed using GraphPad Prism 10. Ordinary One-way ANOVA analysis or unpaired *t* tests were performed to determine statistical significance of the data. Graphs show mean ± SD (n = 3 technical replicates unless specified in figure legend). *\*P < 0.05, **P < 0.01, ***P <0.001* and *\*\*\*\*P <0.0001* were considered statistically significant.

## Abbreviations

EGF: epidermal growth factor
ER: endoplasmic reticulum
F-actin: filamentous actin
FDA: Federal Drug Agency
HBSS: Hanks balanced salt solution
HHD: Hailey Hailey Disease
IGF: insulin like growth factor
PBS: phosphate buffered saline
PFA: paraformaldehyde
3D: Three Dimensional;
TEM: transmission electron microscopy

