## Supplementary material for "Insulin-Regulated Actin Dynamics is Disrupted in a Human Keratinocyte Model of Hailey Hailey Disease": All Supplemental Figures

### Slide 1
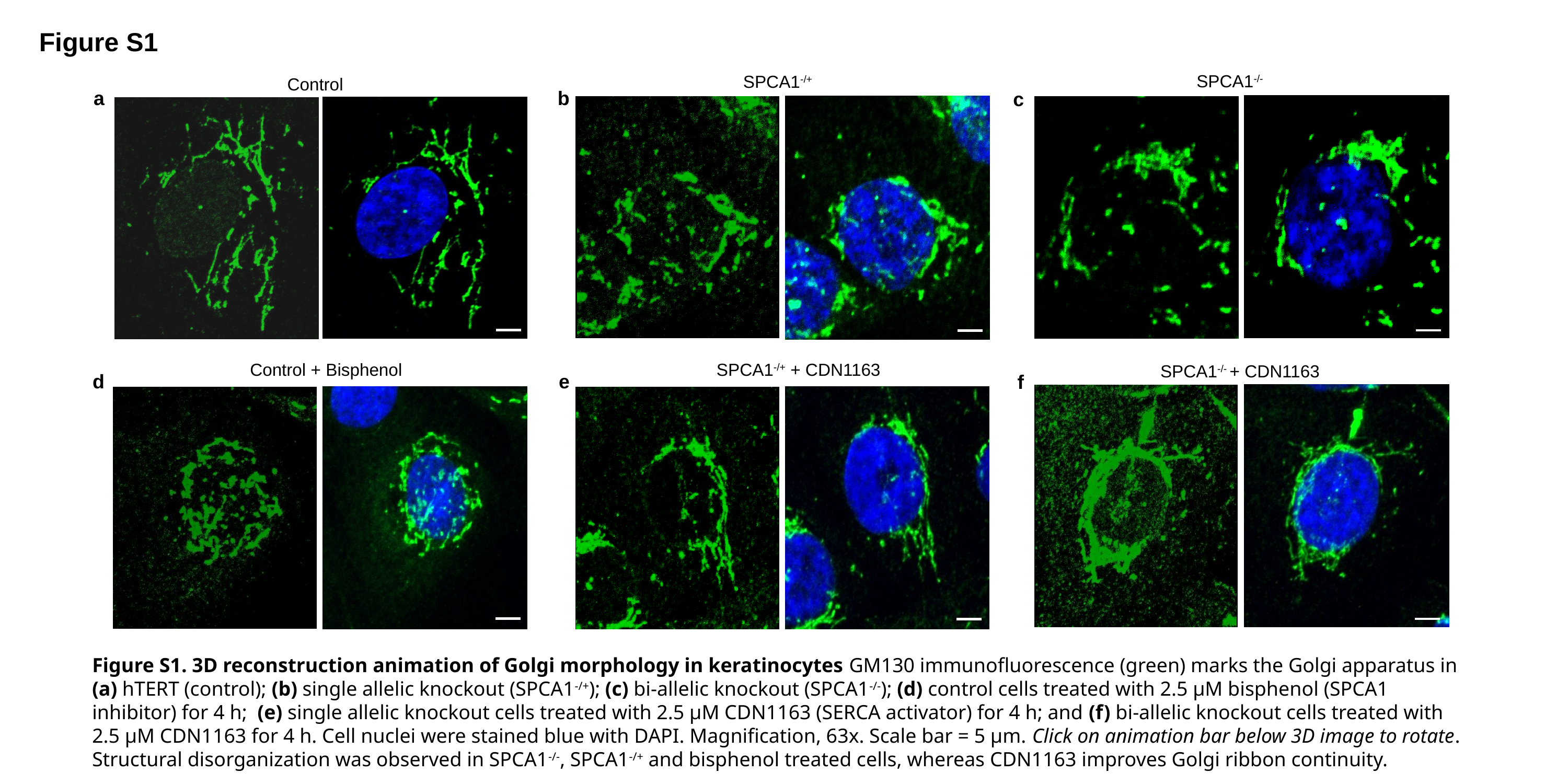

Figure S1
SPCA1^-/-^
SPCA1^-/+^
Control
a
b
c
SPCA1^-/+^ + CDN1163
Control + Bisphenol
SPCA1^-/-^ + CDN1163
d
e
f
Figure S1. 3D reconstruction animation of Golgi morphology in keratinocytes GM130 immunofluorescence (green) marks the Golgi apparatus in (a) hTERT (control); (b) single allelic knockout (SPCA1^-/+^); (c) bi-allelic knockout (SPCA1^-/-^); (d) control cells treated with 2.5 µM bisphenol (SPCA1 inhibitor) for 4 h; (e) single allelic knockout cells treated with 2.5 µM CDN1163 (SERCA activator) for 4 h; and (f) bi-allelic knockout cells treated with 2.5 µM CDN1163 for 4 h. Cell nuclei were stained blue with DAPI. Magnification, 63x. Scale bar = 5 µm. Click on animation bar below 3D image to rotate. Structural disorganization was observed in SPCA1^-/-^, SPCA1^-/+^ and bisphenol treated cells, whereas CDN1163 improves Golgi ribbon continuity.

### Slide 2
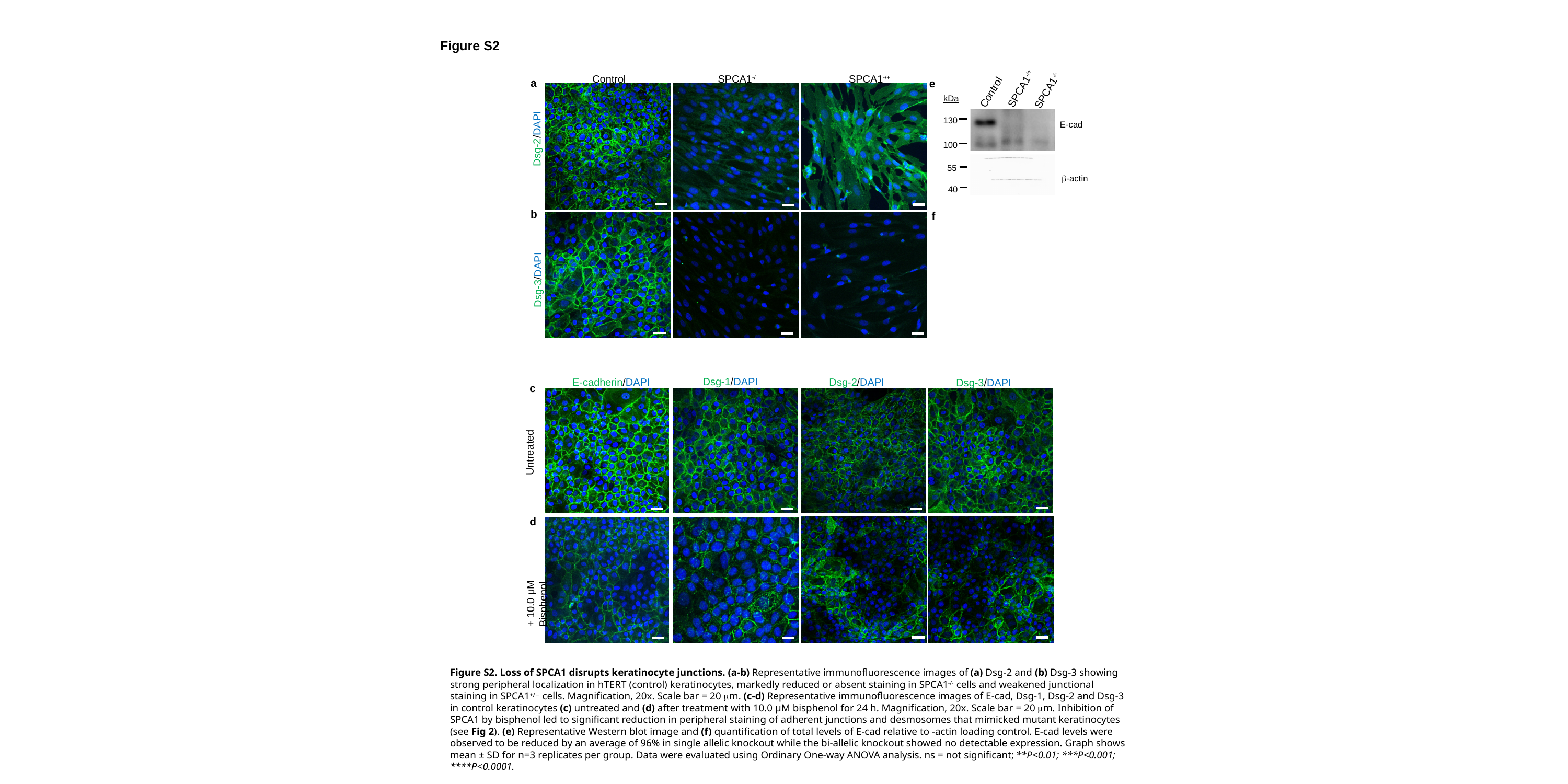

Figure S2
Control
SPCA1^-/-^
SPCA1^-/+^
a
e
SPCA1^-/+^
SPCA1^-/-^
Control
kDa
130
E-cad
Dsg-2/DAPI
100
55
β-actin
40
b
f
Dsg-3/DAPI
Dsg-1/DAPI
Dsg-2/DAPI
E-cadherin/DAPI
Dsg-3/DAPI
c
Untreated
d
+ 10.0 μM Bisphenol

### Slide 3
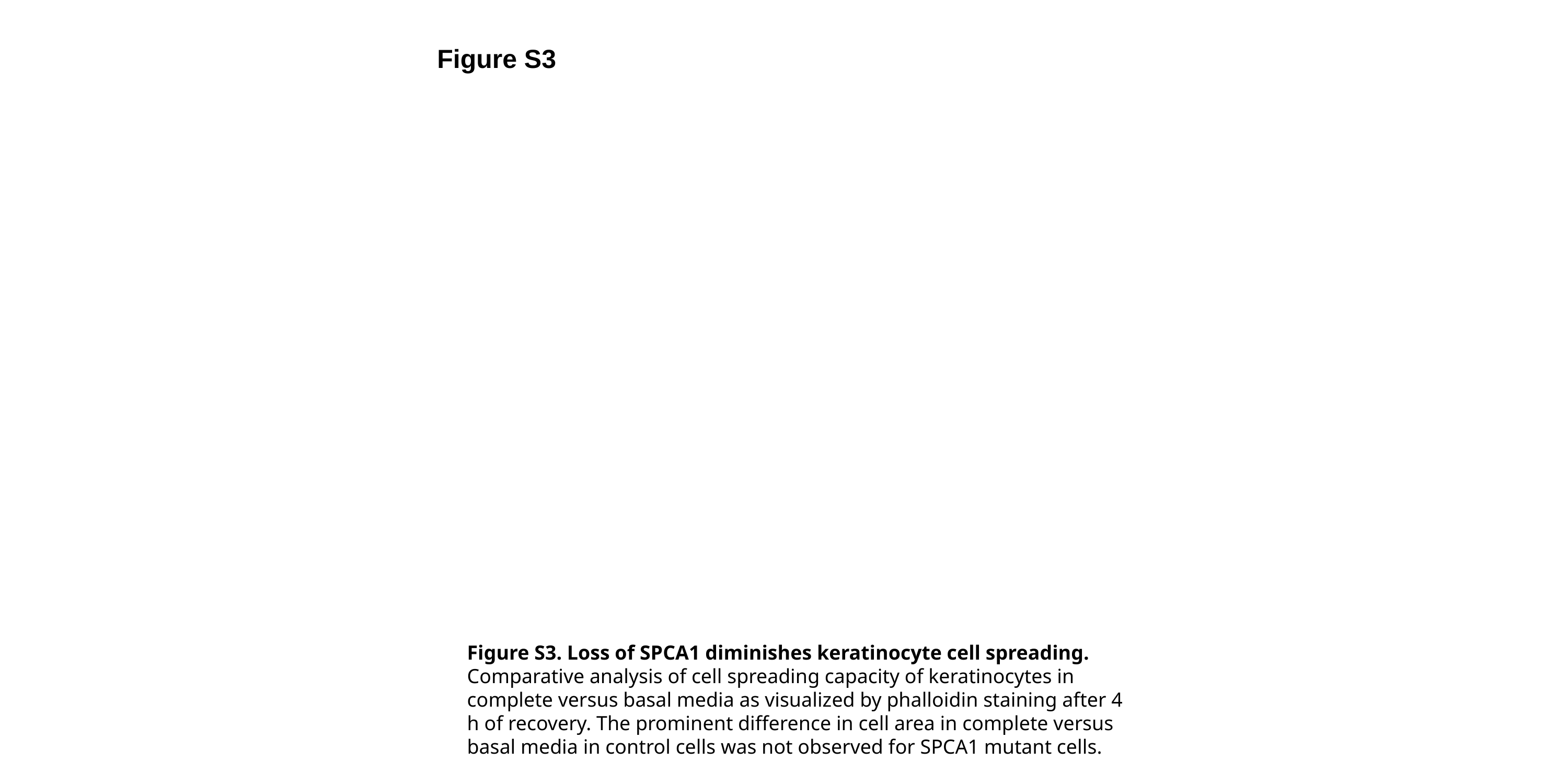

Figure S3
Figure S3. Loss of SPCA1 diminishes keratinocyte cell spreading. Comparative analysis of cell spreading capacity of keratinocytes in complete versus basal media as visualized by phalloidin staining after 4 h of recovery. The prominent difference in cell area in complete versus basal media in control cells was not observed for SPCA1 mutant cells.

### Slide 4
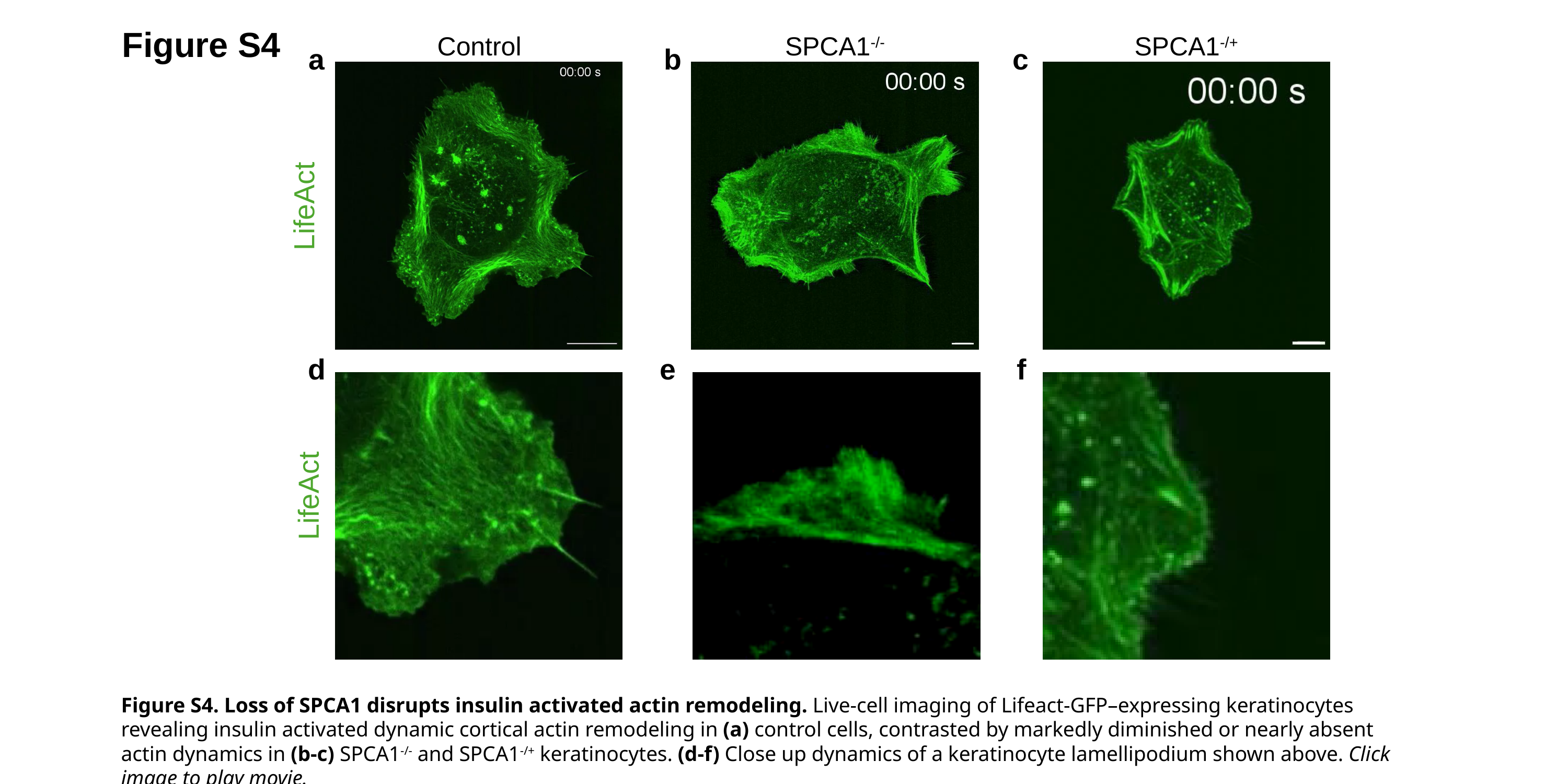

Figure S4
Control
SPCA1^-/-^
SPCA1^-/+^
a
b
c
LifeAct
d
e
f
LifeAct
Figure S4. Loss of SPCA1 disrupts insulin activated actin remodeling. Live-cell imaging of Lifeact-GFP–expressing keratinocytes revealing insulin activated dynamic cortical actin remodeling in (a) control cells, contrasted by markedly diminished or nearly absent actin dynamics in (b-c) SPCA1^-/-^ and SPCA1^-/+^ keratinocytes. (d-f) Close up dynamics of a keratinocyte lamellipodium shown above. Click image to play movie.

### Slide 5
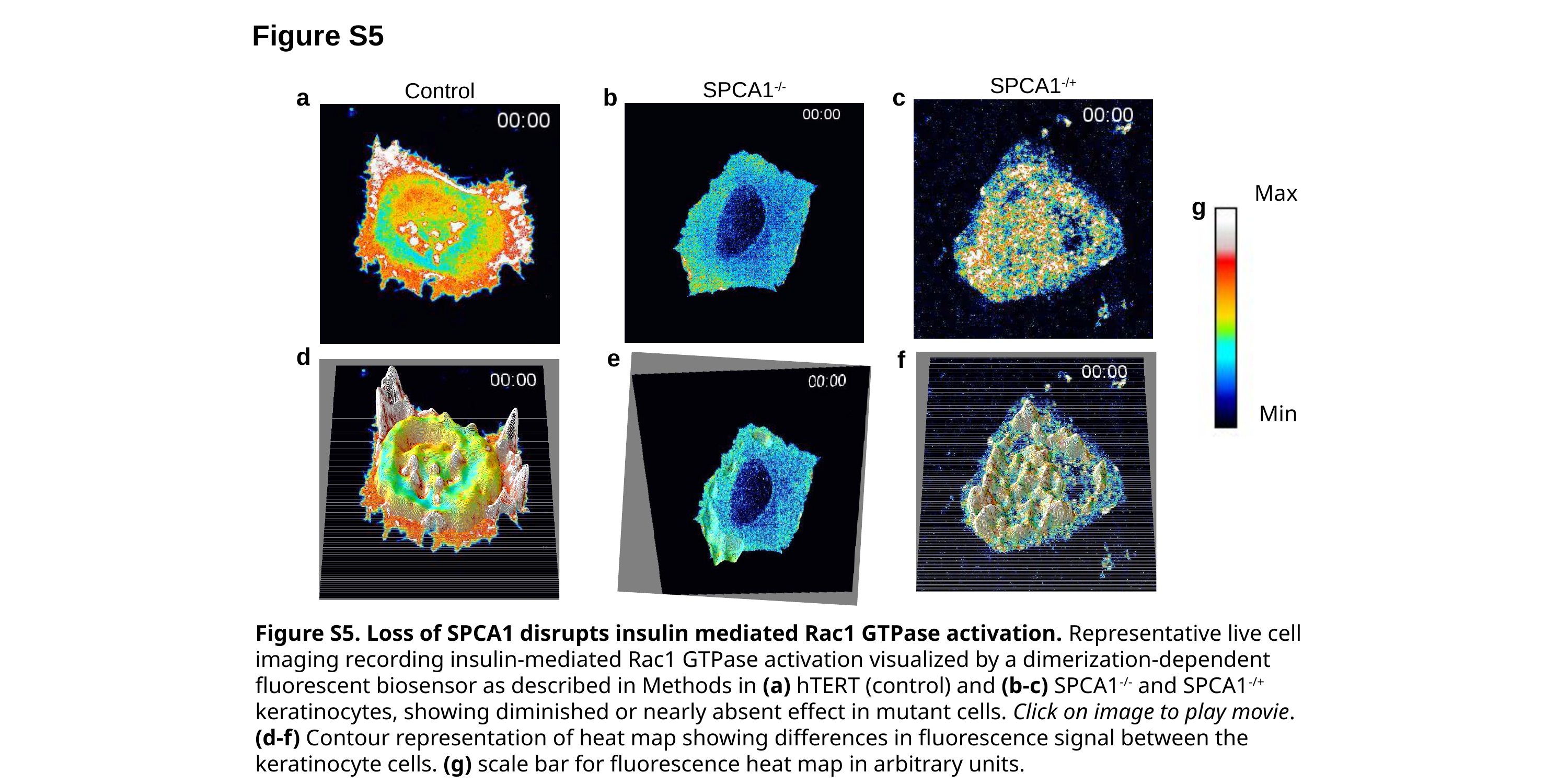

Figure S5
SPCA1^-/+^
SPCA1^-/-^
Control
a
b
c
Max
Min
g
d
e
f
Figure S5. Loss of SPCA1 disrupts insulin mediated Rac1 GTPase activation. Representative live cell imaging recording insulin-mediated Rac1 GTPase activation visualized by a dimerization-dependent fluorescent biosensor as described in Methods in (a) hTERT (control) and (b-c) SPCA1^-/-^ and SPCA1^-/+^ keratinocytes, showing diminished or nearly absent effect in mutant cells. Click on image to play movie. (d-f) Contour representation of heat map showing differences in fluorescence signal between the keratinocyte cells. (g) scale bar for fluorescence heat map in arbitrary units.
